# ARSA: an autonomous research scientist for target nomination in Alzheimer’s disease and related dementias

**DOI:** 10.64898/2026.09.19.752836

**Authors:** Xueyang Li, Jiahui Liu, Jiannan Liu, Timothy I. Richardson, Jeffrey L. Dage, Rebecca C. Klein, Danny Z. Chen, Kun Huang, Jie Zhang, Travis S. Johnson, Yiyu Shi

## Abstract

Expanding therapeutic options for Alzheimer’s disease and related dementias (ADRD) requires biologically grounded targets, yet nomination demands labor-intensive analysis and multidisciplinary evidence synthesis. To address this challenge, we present ARSA, an autonomous research scientist that transforms natural-language research interests and molecular data into prioritized, evidence-grounded target shortlists. ARSA formulates and audits hypotheses, adapts molecular analyses to observed results, and prioritizes targets using cross-cohort evidence and disease-specific knowledge, preserving the evidence and decisions underlying each nomination. Across three complementary evaluations, we show that ARSA generates hypotheses corresponding to subsequent research and identifies credible candidates within and beyond community nomination records. In structured assessment by 14 experts spanning all four technical cores of the Indiana University School of Medicine–Purdue University TREAT-AD Center, every expert assigned higher mean credibility to ARSA-retained candidates than to rejected comparators. ARSA enables systematic, transparent target exploration, opening opportunities to broaden the therapeutic mechanisms investigated in ADRD.

## 1 Introduction

Alzheimer’s disease and related dementias (ADRD) impose a substantial global burden, with an estimated 57 million people living with dementia in 2021 [1]. Despite progress with anti-amyloid treatment in early Alzheimer’s disease (AD) [2], expanding the range of therapeutic mechanisms remains a major priority [3]. Target nomination connects disease biology to therapeutic development by identifying and prioritizing molecular candidates for experimental validation and drug discovery [4]. These upstream decisions direct subsequent research investment [5], and their importance is reflected in the greater clinical success of drug mechanisms supported by human genetic evidence [6]. A more systematic and efficient approach to nominating biologically grounded targets could therefore accelerate early-stage ADRD drug discovery and broaden the mechanisms advanced for therapeutic investigation.

However, developing a defensible target nomination remains a labor-intensive process that requires coordination across multiple disciplines [4]. Researchers must review published studies, formulate mechanistic hypotheses, analyze molecular data, and assemble evidence for a candidate’s disease relevance and therapeutic tractability. Implementing these analyses requires programming and statistical expertise [7], while interpreting their therapeutic implications draws on disease biology, structural biology, pharmacology, and medicinal chemistry [4]. The capacity to investigate new candidates therefore depends not only on access to data, but also on access to specialized expertise and the time required to coordinate these activities. In ADRD, this work is further complicated by the need to reconcile cohort-specific clinical and pathological definitions [8, 9], interpret molecular changes in their cellular and anatomical contexts [10, 11], and weigh discordant transcriptomic and proteomic evidence [12]. Evidence platforms such as Open Targets [13] make relevant information accessible, but investigators must still determine which analyses to perform and how the combined evidence supports a nomination. This dependence on manual analysis planning and evidence synthesis limits the scale at which research interests can be translated into candidates for experimental follow-up. An autonomous framework that connects literature review, hypothesis-directed molecular analysis, and target-specific evidence assessment is therefore needed to reduce this coordination burden and expand systematic ADRD target exploration.

Recent advances in large language model (LLM) agents provide a practical basis for automating these research activities. Biomni [14] dynamically constructs workflows for causal-gene prioritization, drug repurposing, and multimodal analysis, while BioMedAgent [15] coordinates specialized tools for multistep biomedical analyses. CellVoyager [16] generates and executes new analyses of single-cell datasets, and systems such as Co-Scientist [17] and Robin [18] connect hypothesis generation with experimental investigation. Disease-focused efforts have also emerged: Saichandran et al. [19] automate the statistical testing of previously formulated ADRD hypotheses and Text-to-Target [20] integrates literature knowledge with omics evidence for target prioritization. These advances demonstrate the promise of agentic AI for scientific discovery. However, translating an open-ended ADRD research interest into evidence-supported therapeutic target nominations remains a distinct methodological challenge. The scientific objective is not simply to analyze disease data, but to formulate a research hypothesis, investigate its molecular basis, and determine which candidates warrant therapeutic follow-up. This investigation must connect findings across heterogeneous cohorts, resolve their cellular and anatomical contexts, and assess convergent molecular evidence alongside disease relevance and therapeutic tractability [21]. These are interdependent requirements of the target-nomination task, not individual analyses that can be evaluated in isolation. Yet, to the best of our knowledge, no prior work has demonstrated an end-to-end agentic workflow that begins with an open-ended ADRD research interest and carries it through to therapeutic target nomination. Autonomous ADRD target nomination therefore remains an open methodological problem.

To address these challenges, we introduce ARSA (Autonomous Research Scientist for Alzheimer’s Disease and Related Dementias), the first agentic framework designed specifically for ADRD target nomination. Starting from a natural-language research interest and user-provided molecular datasets, ARSA generates literature-grounded hypotheses and autonomously investigates a user-selected hypothesis to produce a prioritized shortlist of candidate therapeutic targets. Its central design principle is to make the evidence required for target nomination guide both analytical planning and candidate selection, connecting cohort-aware molecular analysis to disease relevance and therapeutic tractability while preserving the basis of each nomination. We evaluate ARSA through retrospective correspondence with subsequent literature, concordance with community nominations, and structured expert assessment of target credibility and novelty. The evaluations show that ARSA generates hypotheses corresponding to later research and nominates meaningful candidates both within and beyond existing community records. Fourteen researchers spanning all four technical cores of the Indiana University School of Medicine–Purdue University TREAT-AD Center [4] assessed selected candidates. Notably, retained candidates absent from Agora received a mean within-questionnaire credibility score 1.06 points higher than rejected comparators on a five-point scale, with the same direction of difference in 15 out of 16 questionnaires. Together, these findings demonstrate that ARSA connects open-ended ADRD inquiry to evidence-supported target nominations, including candidates beyond the community registry that domain experts consider credible for further investigation.

## 2 Results

### ARSA Methodology

ARSA is an agentic framework specifically built for ADRD target nomination, translating a natural-language research interest and user-provided molecular data into a prioritized, auditable shortlist of candidate therapeutic targets. The system autonomously retrieves relevant literature, formulates candidate hypotheses, and uses claim-level auditing to guide their refinement. For a hypothesis selected by the user, ARSA plans and executes molecular analyses that account for cohort-specific diagnostic definitions and cell-type annotations. Rather than stopping at disease-associated genes, ARSA makes cross-cohort molecular support, AD-specific evidence, and therapeutic tractability explicit criteria for target nomination. Evidence-centered skills connect these criteria to analytical routines and resource queries, while a defined scoring rubric determines candidate rankings and retention decisions. Each investigation produces a Decision Dossier linking nominations to their supporting evidence, analytical rationale, and limitations, together with a Computational Notebook that re-derives key outputs and checks their consistency with saved records. The overall workflow is illustrated in Fig. 1a, with detailed implementation procedures, model configurations, and prompts provided in the Methods.

**Fig. 1.**
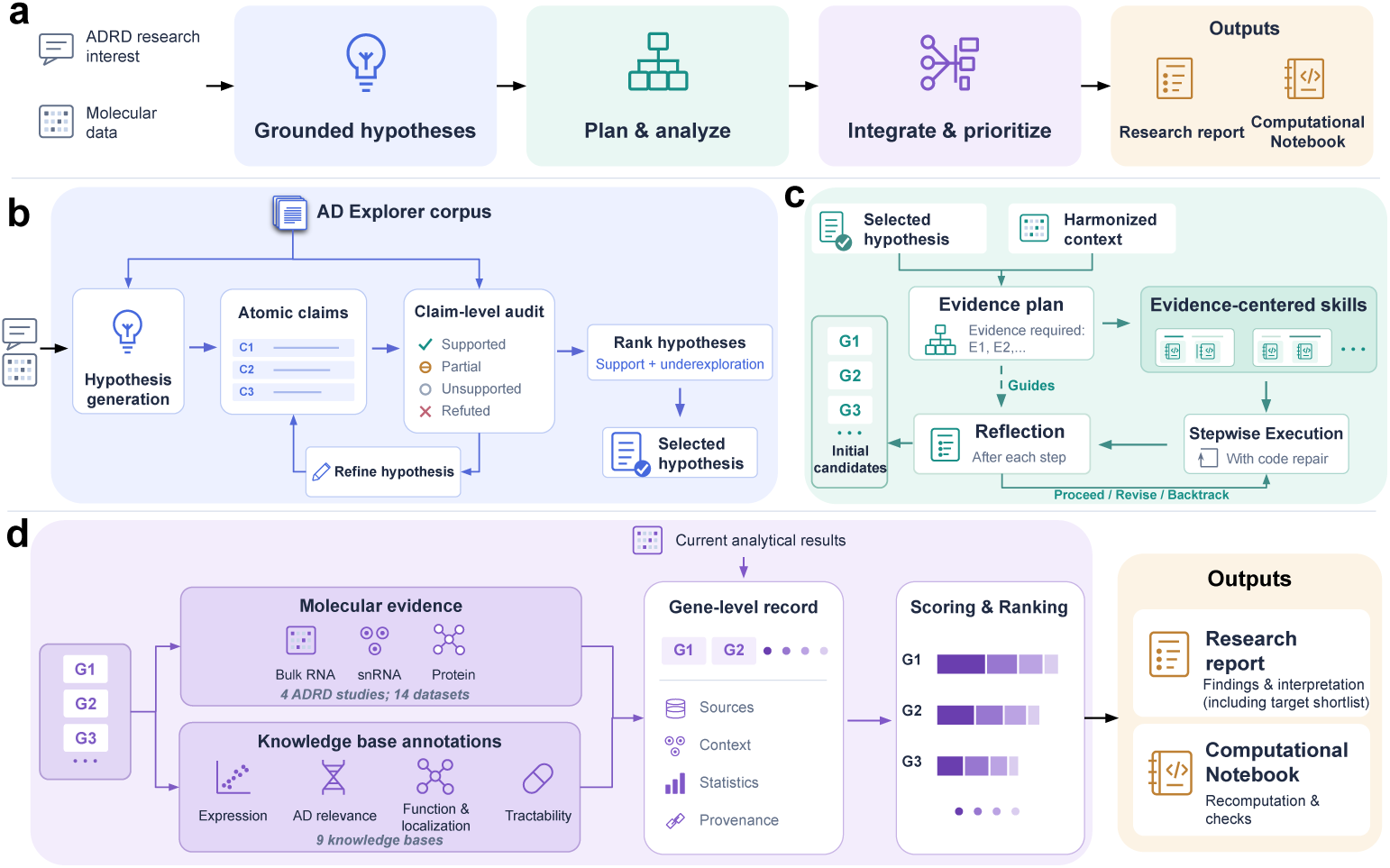
ARSA architecture for autonomous target nomination. **a**, Workflow from an ADRD research interest and molecular data to target nomination and research outputs. **b**, The research interest, available dataset context, and literature from the AD Explorer corpus guide hypothesis generation. Claim-level auditing supports refinement and reassessment, followed by hypothesis ranking and user selection. **c**, The selected hypothesis and harmonized dataset context guide evidence-centered planning and execution. Code repair addresses execution failures, while reflection after each step determines whether to proceed, revise, backtrack, or terminate the investigation (termination branch not shown). Completed analyses yield initial candidate genes. **d**, Current analytical results are integrated with reference evidence from 14 datasets across four ADRD studies: ROSMAP [22–25], the Mount Sinai Brain Bank (MSBB) [26–28], Mayo [29], and SEA-AD [9, 30], together with annotations from nine knowledge bases. Molecular resources include bulk and single-nucleus RNA sequencing (snRNA) and proteomics. Knowledge-base categories comprise expression (GTEx [31]); AD relevance (Agora [32], GWAS Catalog [33], and Open Targets [34]); function and localization (Gene Ontology [35, 36], Reactome [37], and UniProt [38]); and tractability (ChEMBL [39], Pharos [40], and Open Targets). Gene-level evidence records support additive scoring, candidate retention, and ranking. Outputs comprise a research report (Decision Dossier) and a Computational Notebook. The report presents findings and biological interpretation, including the prioritized target shortlist and its supporting evidence, while the notebook supports recomputation and consistency checks. C1–C3, E1–E2, and G1–G3 denote illustrative claims, evidence requirements, and candidate genes, respectively. Stacked bars schematically represent scoring contributions.

### Literature-grounded hypothesis generation

ARSA accepts a natural-language research interest, optionally accompanied by molecular datasets supplied through file uploads or file paths accessible to the system. The research interest defines the scientific objective, while available dataset metadata provide the tissue, cell-type, disease, and modality context for hypothesis formulation. To establish the literature background, ARSA searches an abstract corpus constructed from an AD Explorer [41] source collection of 202,333 PubMed-indexed publications. An LLM uses the retrieved literature and available data context to formulate candidate hypotheses, each accompanied by a literature-based rationale and proposed analytical directions. ARSA then examines the literature basis of each candidate before proceeding to analytical planning. Evidence matching considers genes, brain regions, disease categories, and research topics, allowing retrieval to account for the biological context of the proposed investigation. Rather than treating a relevant publication as support for an entire hypothesis, the system assesses atomic claims within the hypothesis and its rationale. When direct support is insufficient, claim-level assessments guide iterative revision and reassessment, with instructions to retain supported statements, qualify or remove unsupported content, and remove refuted claims (Fig. 1b). Literature therefore informs not only which hypotheses are proposed, but also how their scientific rationale is refined.

To prioritize directions for further investigation, ARSA considers existing support and potential novelty separately. Claim-level support and estimated source breadth characterize the evidence base, while relative sparsity in the literature representation space serves as a proxy for underexploration. Prior therapeutic experience further informs prioritization through a ranking penalty for hypotheses matching historically unsuccessful mechanism classes. The ranked hypotheses are retained with their supporting evidence and scoring contributions for user inspection, and the selected hypothesis guides subsequent analytical planning and execution.

### Hypothesis-directed analytical planning and execution

ARSA translates the selected hypothesis into a molecular investigation by establishing the disease comparison and cellular context of the available data. To accommodate cohort-specific gene identifiers, diagnostic definitions, and cell-type annotations, it uses HGNC-based identifier mapping to link gene-level information across resources, applies cohort-specific diagnostic rules to the available metadata to assign AD and control groups, and maps cell-type labels to a shared vocabulary. With the disease groups and cellular annotations established, the planner selects hypothesis-relevant populations from those represented in the data for AD-versus-control analyses.

Building on this harmonized context, the planner determines which evidence is needed and specifies the skills, analysis parameters, and scientific rationale for obtaining it. Evidence-centered skills organize analytical routines and resource queries by the biological evidence they provide, rather than by data modality alone. For example, the cohort replication skill brings together transcriptomic queries across ADRD cohorts, allowing the planner to request cross-cohort support as a single evidence requirement rather than specify each underlying resource query. Data modality informs the selection of compatible operations within this evidence-centered plan.

ARSA executes the analytical plan stepwise, using predefined tools and falling back to code generation when tool execution fails. Code repair addresses execution errors, while reflection after each step determines whether to proceed, revise the analysis, return to an earlier step, or terminate the investigation (Fig. 1c). Completed analyses provide the basis for initial candidate selection. For differential-expression analyses, gene-level effect sizes and adjusted *P* values guide selection according to the specified criteria. Hypothesis-related focal genes represented in the analysis may also be carried forward when these criteria are not met, with their observed statistics retained and their inclusion explicitly marked. The resulting candidate set undergoes systematic reference-evidence collection, independently of the analytical skills selected during planning.

### Evidence integration and target prioritization

A central challenge in ADRD target nomination is determining which disease-associated genes warrant further investigation as therapeutic targets, rather than merely detecting molecular changes. Transcriptomic and proteomic findings can show distinct or opposing disease associations [12], making it necessary to evaluate candidates against complementary evidence beyond the dataset in which they were identified. For each candidate gene, ARSA integrates the current analytical results with reference molecular evidence from 14 datasets across four ADRD studies and annotations from nine knowledge bases (Fig. 1d). These sources provide complementary information on transcriptomic and proteomic changes, community nominations, AD genetic and disease associations, tissue expression, sub-cellular localization, biological function, and therapeutic tractability. The collected evidence is consolidated into gene-level records that preserve the sources and biological contexts of individual observations, together with available effect estimates and statistical results.

ARSA then applies a user-configurable scoring rubric that combines the collected evidence with hypothesis-related gene information to prioritize candidates. For each candidate, the system records the contribution of each scoring factor and the resulting selection decision. Candidates meeting the specified score threshold are retained for further investigation, with designated concerns flagged for review. Candidates below the threshold are classified as rejected and excluded from the shortlist, while their evidence records and scoring contributions remain available for inspection. The resulting shortlist presents candidate targets alongside the complete set of collected evidence and the scoring contributions underlying their nomination.

### Research reporting and provenance

ADRD target nominations can guide the allocation of experimental resources, making it essential for researchers to examine the evidence, analytical decisions, and uncertainties underlying each recommendation. ARSA therefore presents each investigation as a structured research report, termed the Decision Dossier, with a companion Computational Notebook. The report brings together candidate nominations and gene-level evidence assessments with the research rationale, biological interpretation, and methodological limitations. It also synthesizes findings across candidates and proposes further investigations to address unresolved questions. The underlying evidence records, scoring tables, and intermediate analytical outputs, including differential expression tables and figures, remain available for inspection alongside the report. Furthermore, the Computational Notebook re-derives the key analytical and scoring outputs underlying the nominations, checks them against the saved records, and halts if a consistency check fails. These complementary outputs allow researchers to evaluate the scientific basis of each nomination and determine which findings warrant further analysis or experimental investigation.

### Evaluation of exploratory ADRD research

ARSA addresses an exploratory task: the therapeutic value of a newly nominated gene is often unknown. No exhaustive ground-truth set is therefore available to classify all candidates as correct or incorrect. Resources such as Agora [42] record community nominations, but do not encompass all potentially valuable targets. Absence from such a resource cannot be treated as evidence that a candidate lacks value. Conversely, recovering an entire nomination registry from a given dataset is not a realistic objective, because its entries arise from diverse cohorts, measurements, and experimental systems.

We therefore used three complementary assessments. First, we restricted literature retrieval to pre-cutoff publications and assessed whether the generated hypotheses corresponded to research reported after that cutoff. The literature cutoff was set after the generating model’s documented knowledge cutoff to reduce the risk of training-data contamination. Second, we measured concordance between ARSA-generated targets and community nominations in Agora while withholding Agora-derived nomination information from the workflow. Because registry concordance cannot establish the value of unlisted candidates, we placed particular emphasis on structured expert assessment of target credibility, with perceived novelty evaluated separately. The panel included researchers from all four technical cores of the Indiana University School of Medicine–Purdue University TREAT-AD Center [4], a National Institute on Aging-funded center for AD target assessment and early-stage drug discovery. Their expertise spanned computational biology, structural biology, assay development, and medicinal chemistry.

### Retrospective correspondence with subsequent ADRD literature

We first evaluated the scientific relevance of ARSA-generated hypotheses by examining whether their proposed research questions were investigated in subsequent ADRD publications. As illustrated in Fig. 2a, we conducted 20 ARSA ideation runs with distinct research-interest prompts, requesting five hypotheses per run and obtaining 100 hypotheses in total. All runs used GPT-4o and restricted literature retrieval to publications before January 1, 2025. This cutoff was set 15 months after GPT-4o’s documented knowledge cutoff (October 1, 2023), providing a temporal buffer to reduce the risk of training-data contamination. The evaluation corpus comprised 30,314 publications from 2025 onward. To assess correspondence at this scale without exhaustive manual screening, we used Claude Opus 4.8 as the primary LLM evaluator and GPT-5.5 as a second evaluator. Both used the same post-cutoff corpus, search tools, and assessment rubric but conducted separate literature searches. Further details are provided in Methods.

**Fig. 2.**
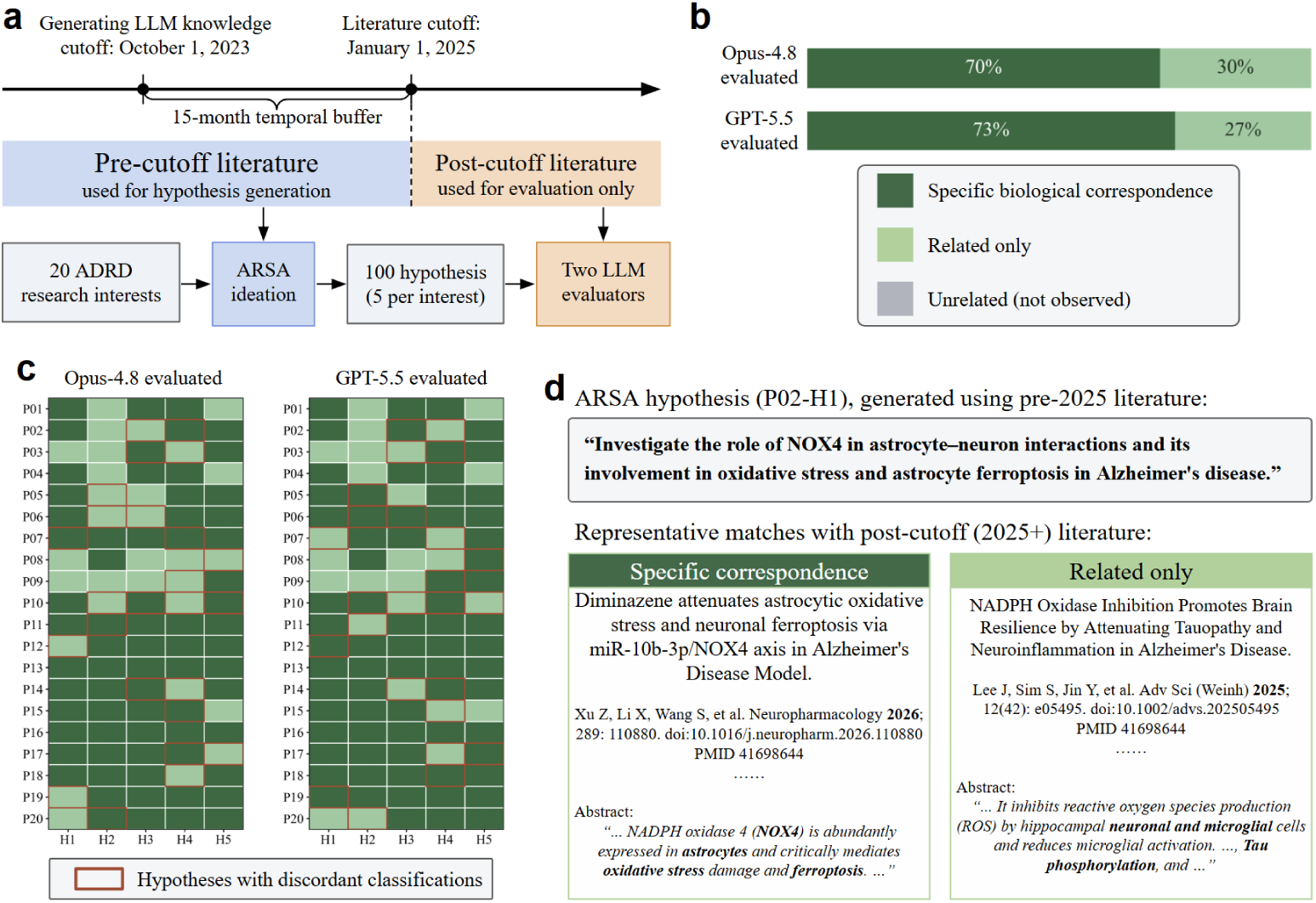
Retrospective correspondence of ARSA-generated hypotheses with subsequent ADRD literature. **a,** Evaluation design. ARSA generated hypotheses using only pre-cutoff literature published before January 1, 2025. The literature cutoff was placed 15 months after the generating model’s documented knowledge cutoff (October 1, 2023) to provide an additional temporal buffer against training-data contamination. Generated hypotheses were then evaluated against post-cutoff ADRD publications dated 2025 or later. **b,** Overall correspondence results for the two evaluators. Topical correspondence was observed for all 100 hypotheses in both evaluations. Specific biological correspondence is a strict subset of topical correspondence; the remaining topically related cases are labeled as related only. **c,** Hypothesis-level correspondence classifications across 20 research interests (P01–P20), each with five generated hypotheses (H1–H5). Dark green indicates specific biological correspondence, light green indicates related-only correspondence, and orange outlines mark hypotheses for which the two evaluators assigned different classifications. Overall agreement on specific biological correspondence was 75% (75/100 hypotheses). **d,** Representative example illustrating the distinction between specific biological correspondence and related-only correspondence for one ARSA-generated hypothesis. Specific biological correspondence required alignment in the genes, mechanism or pathway, cell type, and disease context, whereas related-only correspondence reflected broader topical overlap without meeting all of these criteria.

We applied two nested criteria: relatedness and specific biological correspondence. A hypothesis was classified as related if at least one post-cutoff publication investigated the same biological research area. Specific biological correspondence required a publication to investigate the exact research question posed by the hypothesis: the same entities (allowing synonymous gene names); the same mechanism, pathway, or approach; the same cell type or biological context; and the same disease. Related hypotheses that did not meet this stricter criterion were classified as related only. The assessment captured whether the proposed questions were subsequently investigated, encompassing studies that supported or challenged the hypotheses. Both evaluators classified all 100 hypotheses as related, with no unrelated cases observed. Specific biological correspondence was identified for 70 hypotheses (70%) by Claude Opus 4.8 and 73 (73%) by GPT-5.5 (Fig. 2b).

Claude Opus 4.8 and GPT-5.5 assessed an average of 34.2 and 46.5 candidate publication abstracts per hypothesis, respectively. Their classifications of specific biological correspondence agreed for 75 of the 100 hypotheses (75%), with 25 discordant classifications (Fig. 2c). For example, an ARSA-generated hypothesis on NOX4-mediated astrocyte ferroptosis and astrocyte–neuron interactions in AD was matched to a study examining the miR-10b-3p/NOX4 axis, astrocytic oxidative stress, and neuronal ferroptosis in APP/PS1 mice [43]. A publication classified as related only instead examined pan-NADPH oxidase inhibition, neuronal tau phosphorylation, and microglial inflammatory responses [44] (Fig. 2d). Together, these results support the ability of ARSA’s ideation workflow to formulate hypotheses that correspond to specific biological questions pursued in subsequent ADRD research.

### Concordance with community-nominated targets

We next evaluated whether ARSA’s target nominations agreed with community nominations in Agora [42]. Community nominations provide an external reference for assessing the relevance of ARSA’s outputs to target-directed research, beyond the framework’s own evidence scores. We therefore assessed the proportion of generated candidates represented in the registry, rather than requiring a single dataset to recover nominations derived from diverse research settings. Agora-derived nomination information was withheld from the workflow for this evaluation. We applied the same ten research-interest prompts separately to three postmortem brain single-nucleus RNA-sequencing datasets: GSE138852 from the entorhinal cortex [45], ROSMAP data from the prefrontal cortex [46], and SEA-AD data from the middle temporal gyrus (MTG) [9]. This yielded 30 target-nomination runs, with one shortlist produced for each dataset–prompt combination (Fig. 3a; Methods).

**Fig. 3.**
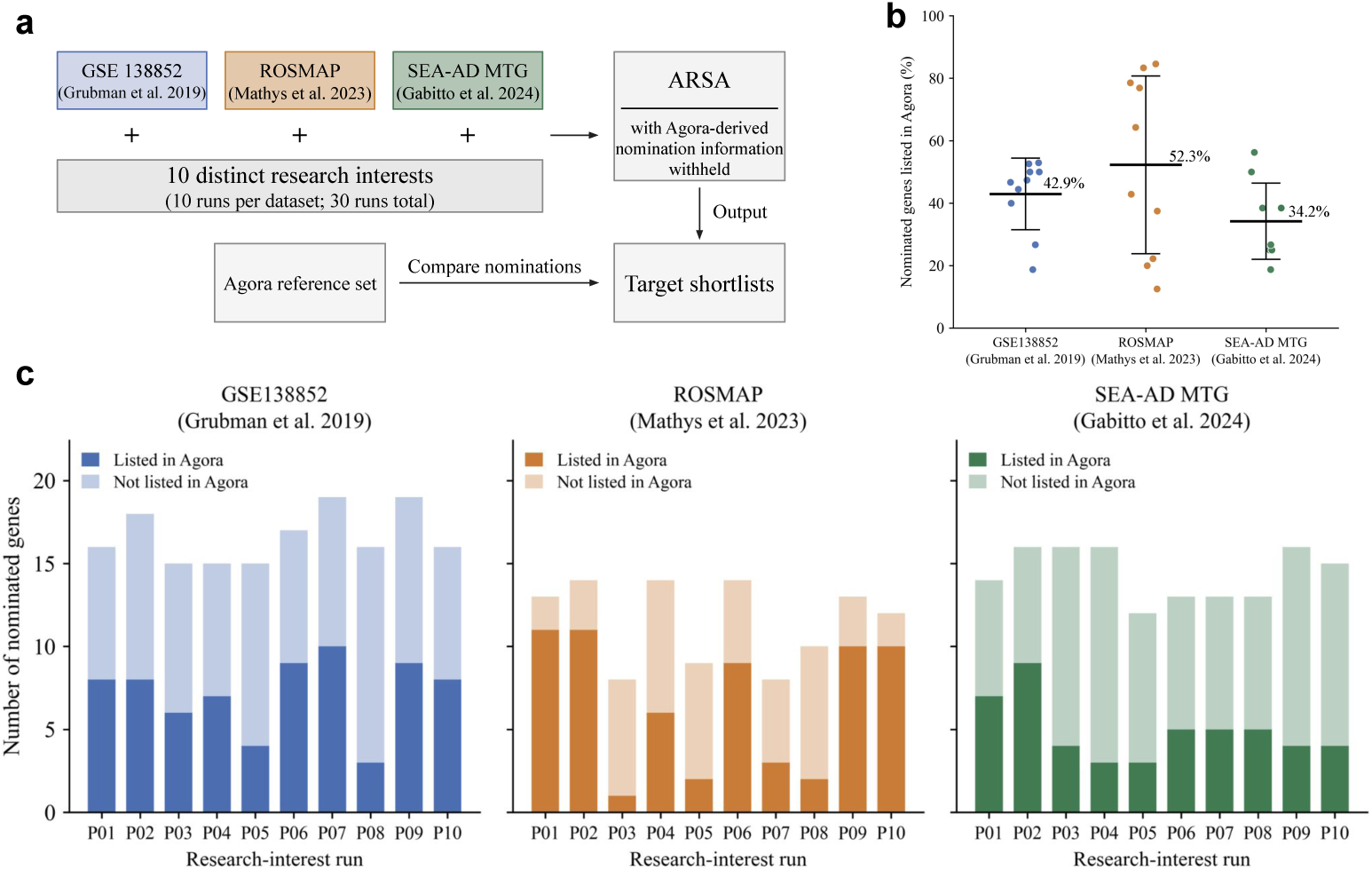
Concordance of ARSA target nominations with community nominations in Agora. **a,** Evaluation design. Ten distinct research-interest prompts were applied separately to GSE138852, ROSMAP, and SEA-AD MTG, yielding 30 runs. Agora-derived nomination information was withheld from the ARSA workflow, and the resulting shortlists were compared with the Agora reference set. **b,** Run-level nomination overlap, calculated as the percentage of retained candidate genes listed in Agora. Each point represents one research-interest run (*n* = 10 per dataset). Black horizontal lines indicate means, and error bars show ±1 s.d. across the ten runs. Annotated percentages indicate the mean overlap for each dataset. The runs differ in research interest and use the same input dataset within each dataset group. **c,** Shortlist size and composition for each run. P01– P10 identify the ten research-interest prompts. Each stacked bar represents one shortlist, with darker segments indicating genes listed in Agora and lighter segments indicating genes not listed in the reference set. Bar heights show the total number of retained candidate genes. Gene counts are reported separately for each run and are not deduplicated across runs.

For each run, nomination overlap was calculated as the proportion of retained candidate genes present in the Agora reference set. As presented in Fig. 3b, mean overlap across the ten runs was 42.9 ± 11.5% for GSE138852, 52.3 ± 28.5% for ROSMAP, and 34.2 ± 12.2% for SEA-AD MTG. All 30 runs produced shortlists containing both Agora-listed genes and candidates absent from the reference set. Thus, ARSA recovered community-nominated candidates without directly using Agora nomination information, while its outputs were not confined to the existing registry.

Shortlists contained 15–19 genes per run for GSE138852, 8–14 for ROSMAP, and 12–16 for SEA-AD MTG (Fig. 3c). Mean recorded runtimes were 5.0, 10.8, and 18.4 min per run for GSE138852, ROSMAP, and SEA-AD MTG, respectively. Across all 30 runs, the mean runtime was 11.4 min, with a median of 8.8 min and a range of 2.8–31.3 min. Timing procedures and computational settings are described in the Methods. In ROSMAP, for example, the microglial-dysfunction run (P01) included the Agora-listed genes *APOE* and *TREM2*, whereas the tau-propagation run (P03) included *CUX2* and *RORB*, neither of which appeared in the reference set. To determine whether nominations beyond the registry warranted further investigation, we next assessed selected candidates through structured expert review of target credibility and perceived novelty.

### Expert assessment of target credibility and novelty

To assess whether ARSA-nominated genes warranted further target-directed investigation, we conducted a structured evaluation with 14 domain experts from the four technical cores of the Indiana University School of Medicine–Purdue University TREAT-AD Center [4]. The panel included faculty investigators and research scientists, with expertise spanning bioinformatics and computational biology, structural biology and biophysics, assay development and high throughput screening, and medicinal chemistry and chemical biology. As shown in Fig. 4a, we randomly sampled 60 genes without replacement within three categories from the gene pool evaluated across the 30 target-nomination runs described above: 20 Agora-listed retained nominations, 30 non-Agora retained nominations, and 10 ARSA-rejected candidates included as comparators. The selected genes were shuffled and distributed across three surveys of 20 genes, each reviewed by at least one expert from every core. Experts assessed gene-level evidence cards using a common rubric, rating target credibility and perceived novelty on separate five-point scales. The panel returned 16 usable questionnaires, yielding 320 gene-level assessments, with four to six ratings per gene for each dimension. Detailed gene-sampling procedures, survey administration, rating criteria, and statistical analyses are provided in the Methods.

**Fig. 4.**
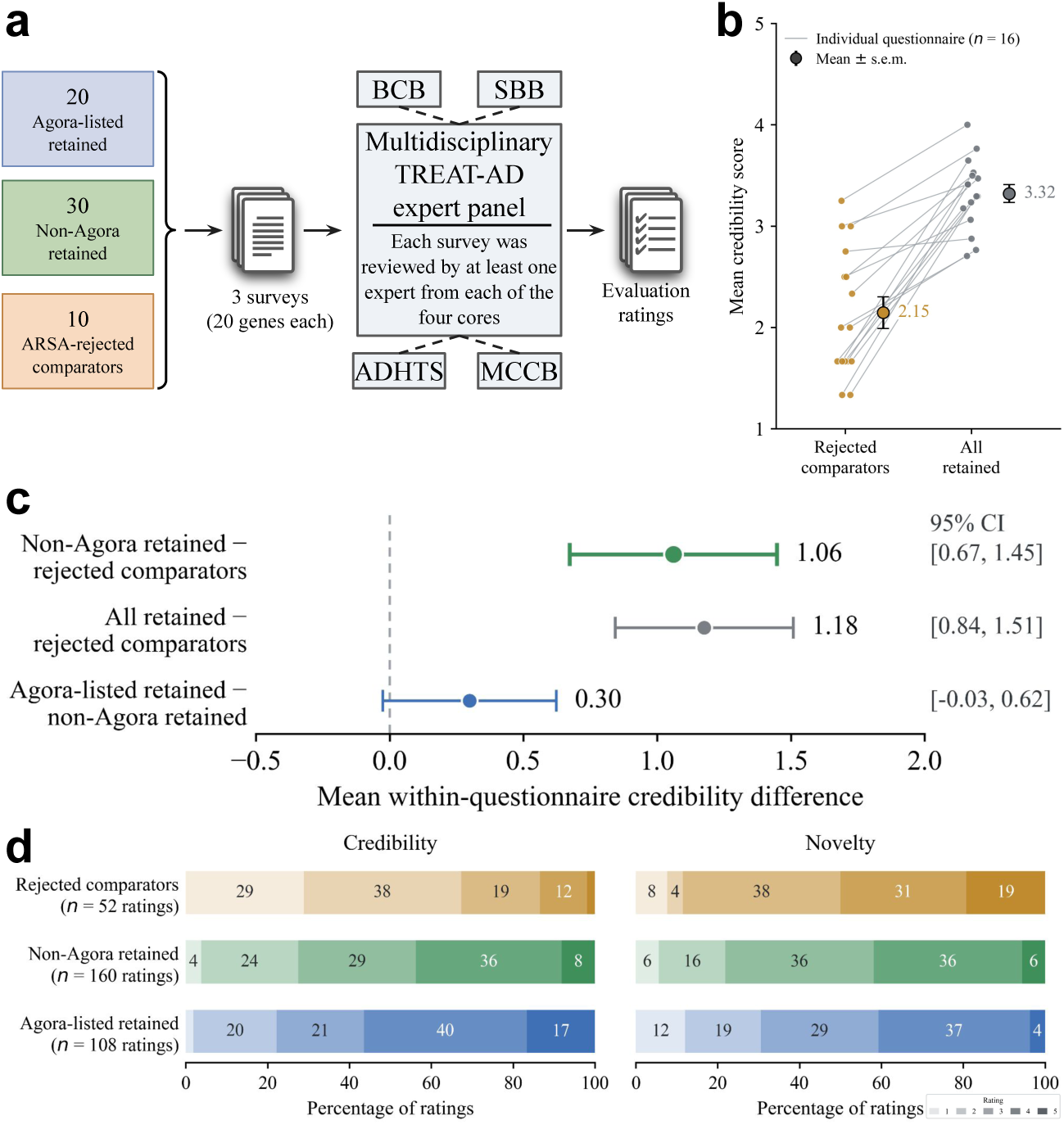
Multidisciplinary expert assessment of ARSA target nominations. **a,** Evaluation design. 60 genes, comprising 20 Agora-listed retained nominations, 30 non-Agora retained nominations, and 10 ARSA-rejected comparators, were shuffled and divided into three surveys of 20 genes each. 14 experts submitted 16 questionnaires. Each survey was reviewed by at least one expert from each of the four TREAT-AD cores: Bioinformatics and Computational Biology Core (BCB), Structural Biology and Biophysics Core (SBB), Assay Development and High Throughput Screening Core (ADHTS), and Medicinal Chemistry and Chemical Biology Core (MCCB). **b,** Mean credibility scores for retained nominations and rejected comparators within each questionnaire. Lines connect means from the same questionnaire; larger circles and error bars indicate the mean ± s.e.m. across the 16 questionnaires. **c,** Mean within-questionnaire credibility differences for the indicated contrasts (*n* = 16 questionnaires). Error bars indicate 95% confidence intervals, and the dashed vertical line marks zero difference. **d,** Distributions of credibility and novelty ratings on five-point scales. Percentages represent shares of individual ratings, not genes. Group sizes were 52 ratings for rejected comparators, 160 for non-Agora retained nominations, and 108 for Agora-listed retained nominations.

ARSA-retained nominations received higher credibility ratings than rejected comparators in every questionnaire. Mean within-questionnaire scores were 3.32 for retained nominations and 2.15 for rejected comparators (Fig. 4b), with a mean paired difference of 1.18 points (95% CI, 0.84–1.51). Crucially, restricting the retained group to non-Agora nominations preserved a difference of 1.06 points (95% CI, 0.67–1.45), again favoring retained candidates in 15 out of 16 questionnaires (Fig. 4c). The separation was therefore not confined to candidates already nominated by the community. An expert-level sensitivity analysis pooling ratings across questionnaires completed by the same expert yielded similar mean credibility differences relative to rejected comparators: 1.17 points (95% CI, 0.80–1.54) for all retained nominations and 1.04 points (95% CI, 0.61–1.46) for non-Agora retained nominations. The all-retained contrast was positive for all 14 experts (exact two-sided sign test, *P* = 1.2 × 10*^−^*^4^), and the non-Agora contrast for 13 of 14 experts (*P* = 1.8 × 10^−3^). The complete rating distributions likewise showed a shift toward higher credibility for retained candidates (Fig. 4d). These results demonstrate consistent agreement between ARSA’s retention decisions and expert judgments, including for nominations outside the reference registry.

At the gene level, 24 of the 30 non-Agora nominations had mean credibility scores of at least 3.0, 10 scored at least 3.5, and two scored at least 4.0 (Fig. 5f). Joint assessment of credibility and novelty revealed patterns not captured by registry membership alone (Fig. 5a). *APOE*, *TREM2*, and *BACE1* received high credibility but low novelty scores, consistent with their status as extensively studied AD-related genes [47–49]. Notably, *BACE1* was absent from the Agora reference set despite the clinical evaluation of BACE1 inhibition as an AD therapeutic strategy [50, 51]. Conversely, the Agora-listed candidate *SLC38A2* received mean scores of 4.00 for both credibility and novelty, with only five gene-linked abstracts in the ADRD corpus. The Y-linked noncoding RNA *TTTY14*, included as a rejected comparator, received low credibility despite relatively high novelty. Thus, absence from Agora did not preclude high credibility, inclusion did not preclude perceived novelty, and novelty alone did not imply expert endorsement.

**Fig. 5.**
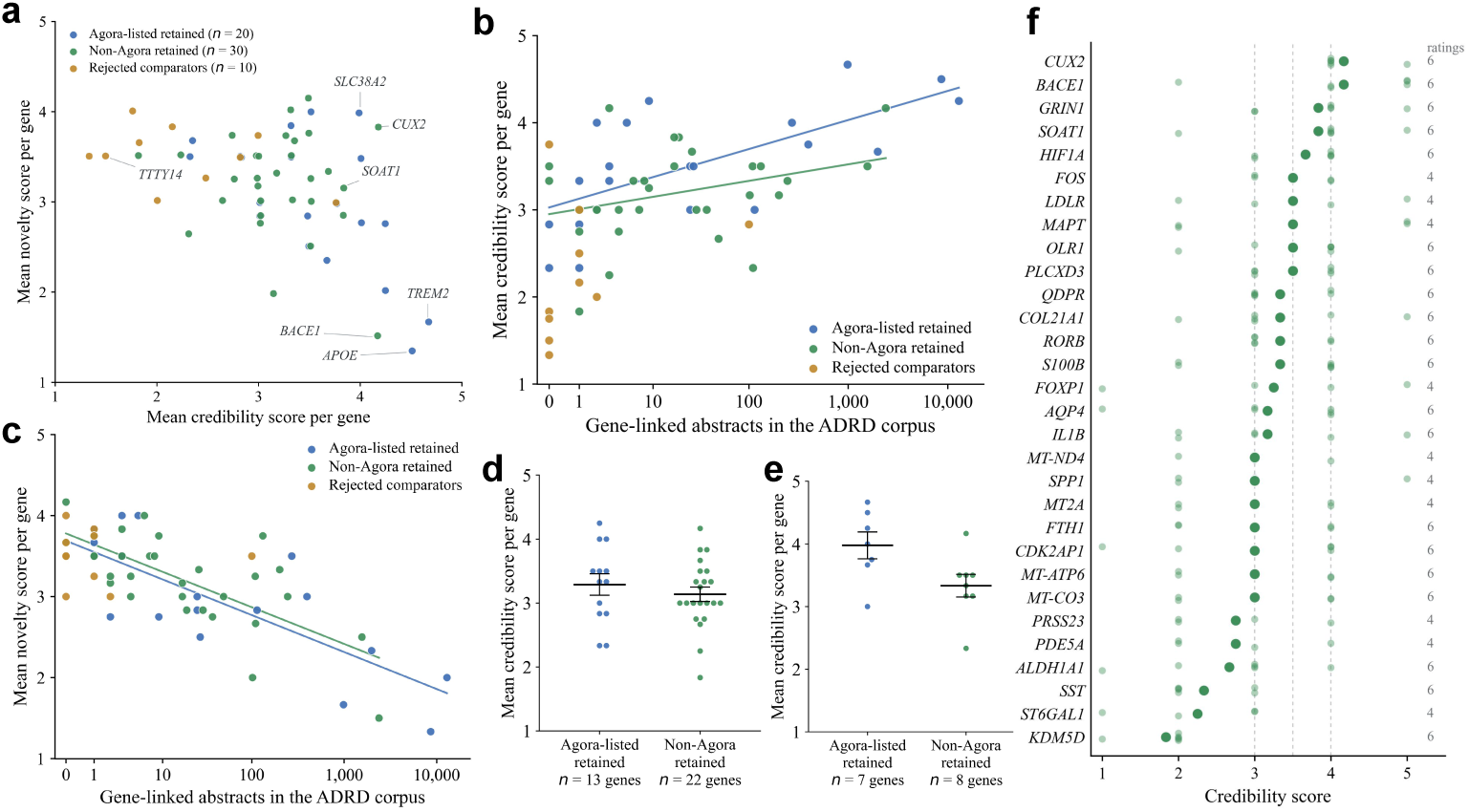
Gene-level credibility, novelty, and literature coverage of evaluated candidates. **a,** Mean credibility and novelty scores for all 60 evaluated genes. Blue, green, and ochre indicate Agora-listed retained nominations (*n* = 20), non-Agora retained nominations (*n* = 30), and rejected comparators (*n* = 10), respectively. Selected genes are labeled. **b,c,** Relationships between literature coverage and gene-level mean credibility (**b**) or novelty (**c**). Literature coverage is measured by the number of gene-linked abstracts in the ADRD corpus. All 60 genes are shown, with separate fitted lines for Agora-listed and non-Agora retained nominations; rejected comparators are displayed but are not included in either fit. Literature-counting and fitting procedures are described in the Methods. **d,e,** Gene-level mean credibility stratified by lower (**d**, fewer than 100 gene-linked abstracts) or higher (**e**, at least 100 gene-linked abstracts) literature coverage. The lower-coverage groups contain 13 Agora-listed and 22 non-Agora retained genes; the higher-coverage groups contain 7 and 8 genes, respectively. Individual points represent genes. Black horizontal lines indicate group means, and error bars show ± s.e.m. across genes. **f,** Credibility scores for all 30 non-Agora retained nominations, ordered by decreasing gene-level mean. Small light-green points indicate individual ratings, and larger dark-green points indicate gene-level means. Numbers at right give the number of ratings for each gene.

We next examined how credibility and novelty ratings varied with the extent of prior ADRD literature on each gene. Fitted credibility scores increased more steeply with literature coverage among Agora-listed nominations than among non-Agora nominations (Fig. 5b). Novelty declined with increasing coverage in both groups, with similar fitted slopes (Fig. 5c), linking perceived novelty to prior research coverage within each registry group. In an exploratory stratified analysis, genes with fewer than 100 linked abstracts had mean credibility scores of 3.29±0.17 for Agora-listed nominations and 3.14±0.11 for non-Agora nominations. Among genes with at least 100 linked abstracts, the corresponding scores were 3.98 ± 0.21 and 3.33 ± 0.18 (mean ± s.e.m. across genes; Fig. 5d,e). Non-Agora nominations in the lower-coverage stratum therefore received mean expert-rated credibility close to that of registry-listed candidates in the same stratum, whereas a larger difference was observed among higher-coverage genes.

## 3 Discussion

ARSA is an autonomous agentic framework for end-to-end ADRD target nomination that transforms a natural-language research interest and user-provided molecular data into a prioritized, evidence-grounded shortlist of candidate therapeutic targets. The contribution of this work is to establish and evaluate a unified approach to autonomous hypothesis-to-target investigation in ADRD, encompassing the formulation of research hypotheses, investigation of a selected hypothesis, and prioritization of candidates for therapeutic follow-up. Rather than requiring researchers to plan and coordinate these stages separately, ARSA connects them within a single research process. Its evidence-centered design provides the organizing principle: the selected hypothesis directs molecular analysis, cohort-specific definitions establish the biological context, and evidence across cohorts and biomedical resources informs candidate selection. Our evaluations show that ARSA generates hypotheses corresponding to subsequent research and produces nominations that overlap community records while also identifying candidates outside those records that domain experts consider credible. Together, these findings demonstrate a practical approach to autonomous hypothesis-to-target investigation, in which the research question, analytical evidence, and nomination rationale remain connected throughout the process.

The three evaluations provide complementary evidence of research value: correspondence with literature published beyond the generating model’s knowledge cutoff connects generated hypotheses to questions subsequently pursued by other investigators, registry concordance situates nominations within existing community research, and expert assessment examines whether candidates warrant further investigation. For exploratory target nomination, concordance with an existing registry provides a reference for agreement with prior research, rather than an objective to maximize in isolation. Its interpretation must be complemented by assessment of the credibility and research value of candidates outside the registry. Our expert evaluation provides this additional evidence: non-Agora retained nominations received higher credibility ratings than rejected comparators in 15 out of 16 questionnaires (mean within-questionnaire difference, 1.06 points; exact two-sided sign test, *P* = 5.2×10^−4^). Thus, expert support for ARSA’s nominations extended beyond recovery of existing registry entries. The contrasting profiles of *BACE1* and *SLC38A2* further distinguish registry membership from perceived credibility and novelty. *BACE1* was absent from the reference set yet received high credibility and low novelty ratings, whereas registry-listed *SLC38A2* was rated highly on both dimensions. Moreover, novelty ratings declined with literature coverage in both registry groups, indicating that perceived novelty was associated with prior research attention rather than registry status alone. Taken together, these findings show that neither registry membership nor literature coverage alone captures the research value of a candidate. Evaluation should therefore recognize both concordance with prior research and the identification of credible candidates beyond existing nomination records.

Although ARSA automates computational investigation through target nomination and provides the evidence and analytical basis for each candidate, selecting candidates for experimental follow-up, designing interventions, and validating their biological effects remain researcher-led activities. The evaluated nomination runs had a mean recorded runtime of 11.4 min (median, 8.8 min), supporting rapid execution of the computational workflow. However, the time saved relative to comparable investigator-led workflows and any gains in downstream experimental productivity remain to be quantified. At the analytical level, ARSA uses conventional bioinformatics procedures for intermediate operations. The present study focuses on the research value of the integrated hypothesis-to-target workflow. Accordingly, our evaluations assessed the generated hypotheses and target nominations without separately bench-marking intermediate operations such as quality control and differential-expression analysis. Future evaluation could compare these intermediate outputs with expert-curated reference analyses to assess the execution of individual analytical steps. Complementary prospective comparisons of investigator-led and ARSA-based nomination workflows could measure researcher time, candidate-selection decisions, and the experimental resources required to advance nominated targets.

In summary, ARSA establishes an end-to-end agentic framework for ADRD target nomination, transforming natural-language research interests and molecular datasets into prioritized, evidence-grounded target shortlists. The framework pairs its nominations with supporting evidence and executable notebooks, enabling researchers to inspect their scientific rationale and reproduce key computations. We hope that ARSA will lower the technical barriers to systematic target exploration and reduce the time required to translate biological questions into candidates for experimental investigation. By making evidence-driven target nomination more accessible, ARSA could enable broader investigation of underexplored disease mechanisms and accelerate early-stage drug discovery for ADRD.

## 4 Methods

### ARSA implementation

ARSA accepts a natural-language research interest, optionally accompanied by molecular datasets and associated metadata, and uses an ADRD literature corpus and biomedical reference resources to support the investigation. When no molecular data are supplied, ARSA operates in an ideation-only mode, generating and ranking hypotheses without proceeding to molecular analysis. When data are available, the user selects a generated hypothesis for downstream molecular investigation and target assessment (Fig. 1a). The framework is implemented in Python, with expression data and associated annotations represented using AnnData [52]. Language-model calls are configured by processing stage and connected to analytical procedures, evidence queries, and target-scoring rules.

#### Literature-grounded hypothesis generation

The literature corpus is constructed from an AD Explorer [41] source collection of 202,333 PubMed-indexed publications. Background retrieval uses BM25 queries formed from the research interest and available tissue, cell-type, disease, and modality metadata. The evaluated ideation configuration used gpt-4o with a temperature of 0 to identify research directions from this context and generate candidate hypotheses. Each candidate contains a hypothesis statement, a literature-based rationale, and proposed analyses.

Each hypothesis and its rationale are subsequently decomposed into atomic claims for separate evidence assessment (Fig. 1b). To support claim-specific literature matching, hypotheses and publications are annotated with shared contextual fields: genes, brain regions, disease categories, and research topics. The classifier’s encoder also supplies normalized text embeddings for semantic retrieval. Claim-specific matching combines keyword and embedding retrieval, reranking, and weighted overlap between contextual fields.

The audit uses publication titles and abstracts to assess support for individual claims. It retains publication-level judgments and assigns each claim to one of four categories: supported, partially supported, unsupported, or refuted. A claim is classified as supported when at least one retrieved publication is judged to establish it directly. The system performs revisions for the failed hypothesis guided by the claim-level assessments. Revision instructions retain supported content, adjust or remove unsupported statements, and remove refuted claims. The revised hypothesis undergoes renewed claim decomposition and auditing, after which the support fraction is recalculated. Generation, auditing, and revision use the same configured model; claim auditing also uses a temperature of 0. In addition, underexploration is derived from the local density of the hypothesis in the learned representation space relative to the corpus. Source breadth *B_h_* is also derived from cohort mentions and last-author identifiers in publications judged to provide full or partial support. The historical-mechanism penalty *H_h_* is applied when previously unsuccessful therapeutic mechanisms are found. This design balances literature-grounded support with exploratory opportunity, while accounting for source breadth and prior therapeutic experience. Assessing underexploration after generation allows hypotheses to emerge from the research interest and available data context without restricting ideation to predefined sparse regions of the literature. The audit threshold guides refinement rather than exclusion, allowing candidates with different levels of retrieved support to remain available for comparison. Candidates are ranked by decreasing *R_h_* and presented with their audit records and individual scoring contributions, enabling users to examine both the evidential basis and exploratory rationale of each candidate before selecting a hypothesis for molecular investigation.

#### Hypothesis-directed analytical planning and execution

ADRD cohorts encode disease status and cellular identity using study-specific conventions, while gene identifiers can differ across datasets and reference resources. ARSA therefore applies harmonization skills to establish explicit disease comparisons and a shared cellular vocabulary before hypothesis-directed analytical planning (Fig. 1c), making study-specific definitions explicit inputs to the investigation, rather than requiring the planner to infer biological equivalence from diagnostic codes, cell-type labels, or gene names alone. The resulting context enables the planner to identify hypothesis-relevant populations within defined disease groups and connect their molecular findings to corresponding reference evidence.

Then, the planner receives the selected hypothesis along with dataset context. It uses this context to determine the evidence requirements of the investigation and select compatible analytical skills. Each skill description specifies the evidence produced, supported inputs, and adjustable parameters, linking a scientific evidence requirement to an executable operation. The LLM generates a structured plan containing the selected skills, their scientific rationales, and execution parameters. Step-specific objectives, success criteria, and any specified methodological checks provide the reference for subsequent reflection. Analytical and initial nomination parameters can be adjusted through the supported planning interface. The selected skills are first executed through their predefined Python implementations in dependency order. If a tool fails during execution, ARSA invokes gpt-5.1-codex to generate Python code for the failed analytical step and executes the resulting code. Generated code undergoes a syntax check before execution and an output-presence check afterward. Failed executions trigger code repair, in which the code-generation model revises the script using the original code and related error messages. The repair prompt directs the model to inspect actual data fields and function signatures rather than guess them. Each local execution–repair loop permits up to five attempts, with the error history retained for subsequent reflection-based revision or backtracking. If all attempts fail, the step is recorded as unsuccessful and passed, together with its error history, to the reflection stage before further execution.

Beyond local code repair, ARSA uses stepwise reflection to adapt the investigation to observed results. After every analytical step, whether execution succeeds or fails, the reasoning model assesses the outputs against the step’s scientific objective, success criteria, and specified methodological checks. Its context includes execution status, captured output, newly created result entries and artifacts, summaries of previous and remaining steps, and error and backtracking histories. The model returns a structured decision to `proceed`, `revise`, `backtrack`, or `abort`. For successfully executed steps, revision regenerates the code to address identified analytical errors, including indicated parameter adjustments, and reruns the step, with up to two revisions. Backtracking restores the saved working state before an eligible earlier step and resumes execution from that point; termination preserves the findings obtained before stopping. This execution–reflection loop makes subsequent actions responsive to both computational outcomes and their adequacy for the planned investigation, rather than advancing through a fixed sequence regardless of the results.

Intermediate analytical outputs and execution parameters are retained as part of the investigation record, together with reflection decisions and error histories. These records support subsequent execution decisions and enable users to inspect the analysis and reproduce key computations. Completed analyses provide the basis for selecting an initial candidate-gene set under the configured nomination rules. For differential-expression results, these rules apply the specified effect-size and significance criteria. Hypothesis-focal genes may also be carried forward, with their observed results and reasons for inclusion recorded separately. The resulting candidate set defines the scope of systematic reference-evidence collection, independently of the planner’s selection of analytical skills. By preserving the connection between analytical outputs, execution decisions, and candidate selection, ARSA makes the derivation of each candidate available for scrutiny. This transparency allows researchers to examine the supporting evidence and unresolved questions before committing resources to experimental follow-up.

#### Evidence integration and target prioritization

For each gene in the initial candidate set, ARSA supplements the current analytical results with systematically retrieved reference evidence (Fig. 1d). The reference evidence layer comprises 14 datasets from four ADRD studies: ROSMAP [22–25], the Mount Sinai Brain Bank (MSBB) [26–28], Mayo [29], and SEA-AD [9, 30]. These datasets encompass bulk and single-nucleus RNA sequencing and tandem-mass-tag proteomics. To avoid repeatedly processing these large datasets during individual investigations, molecular comparisons are precomputed using cohort-specific diagnostic definitions and neuropathological criteria and stored with their analytical settings and provenance. During evidence collection, ARSA first retrieves cached results appropriate to the required comparison. When the available cache does not cover that comparison, the evidence stage requests additional computation using predefined analytical tools, with code generation as a fallback if tool execution fails. Newly computed results are stored as additional cache entries without overwriting existing records, preserving previous analyses and allowing subsequent investigations to reuse the results. Using the harmonized gene identifiers, ARSA also queries 9 reference knowledge bases for community target nominations, AD genetic and disease associations, tissue expression, subcellular localization, biological function, and therapeutic tractability. For each candidate gene, these results are stored alongside the retrieved or newly computed molecular comparisons. Each observation retains its source, relevant biological context, and available effect estimates and statistical results, together with a record of the query or analysis from which it was obtained. The resulting gene-level evidence records provide the inputs to target scoring while preserving the individual observations underlying each candidate’s assessment. To prioritize candidates, ARSA applies a rubric to the assembled molecular evidence and knowledge-base annotations. Individual contributions, total scores, and retention decisions are saved alongside the collected evidence. This separation between observations and scoring rules allows users to inspect the biological basis of each candidate and the explicit criteria that determine its priority.

#### Research reporting and computational provenance

To make target nominations accessible to scientific scrutiny beyond the original agent session, ARSA assembles a source-linked research report, termed the Decision Dossier, from the saved hypothesis, analytical plan, molecular results, gene-level evidence records, and scoring tables. The report presents the prioritized target shortlist alongside candidate-level evidence summaries and scoring contributions. These results are drawn from saved structured records and supplemented with LLM-generated biological interpretation, methodological limitations, and proposed follow-up investigations. Source records and paths to intermediate tables and figures accompany the report, allowing researchers to examine the observations and analytical decisions underlying each nomination rather than relying on the narrative or final ranking alone.

A companion Computational Notebook provides executable verification of the saved research outputs. Generated from the saved workspace, it reloads differential-expression tables, reconstructs recorded query summaries, and checks statistical tables, evidence-source coverage, scoring contributions, and agreement between recorded retention decisions and saved target lists. Failed checks raise errors during execution, exposing inconsistencies in the checked records. These checks operate on the saved outputs of the original investigation. Together, the Decision Dossier and Computational Notebook connect scientific interpretation with record-level verification: researchers can inspect why a candidate was nominated and check the consistency of the evidence and decisions supporting its inclusion. This design provides a traceable basis for evaluating nominations, examining unresolved questions, and selecting candidates for further experimental investigation.

### Evaluation procedures

#### Retrospective hypothesis evaluation

We designed 20 research-interest prompts spanning target nomination, drug repurposing, biomarker discovery, risk-factor investigation, and mechanistic investigation. The prompts were based on general ADRD research interests rather than post-cutoff publications. Each finalized prompt was submitted to ARSA in ideation-only mode using gpt-4o, without molecular data or dataset metadata, requesting five hypotheses. Literature retrieval was restricted to publications before January 1, 2025, and live external retrieval was disabled. All five hypotheses returned for each finalized prompt were evaluated without manual editing, selection, or replacement. The evaluation corpus comprised 30,314 post-cutoff publications collected using disease-level queries independent of the research-interest prompts. Evaluators received hypothesis titles, rationales, and key genes, with internal fingerprints and BM25 scores omitted. Claude Opus 4.8 and GPT-5.5 independently searched the corpus using the same search tools and assessment rubric. Relatedness and specific biological correspondence were each recorded when at least one retrieved publication satisfied the respective criterion. Correspondence rates were calculated over all 100 hypotheses, and inter-evaluator agreement was the proportion receiving the same classification for specific biological correspondence.

#### Agora-withheld nomination evaluation

We designed ten dataset-conditioned target-nomination prompts, distinct from those used in the retrospective hypothesis evaluation, covering cell types, protein pathologies, and biological pathways or processes. The same prompts were applied separately to GSE138852 [45], ROSMAP [46], and SEA-AD MTG [9], yielding 30 runs. Each run generated three candidate hypotheses under the default setting, and the first returned hypothesis was selected automatically without manual screening. Agora queries were disabled during evidence collection, and the Agora-nomination contribution was excluded from target scoring. All other settings followed the defaults described above. The resulting shortlists were compared with an Agora [42] snapshot dated March 4, 2026, containing 955 unique genes. For each run, nomination overlap was calculated as the percentage of retained candidate genes present in the reference set. Dataset-level summaries report the mean and standard deviation across the ten runs. Elapsed runtime was measured from the start of ideation to completion of report generation.

#### Expert assessment and analysis

Standardized image-based evidence cards were generated from structured records for the selected genes and presented through Google Forms. Each card summarized genetic, transcriptomic, proteomic, tractability, expression, and functional-annotation evidence. To reduce anchoring on prior nominations or system judgments, Agora nomination information, ARSA scores, and retained/rejected labels were withheld; LLM-generated summaries were also excluded. Gene order was shuffled separately for each survey. Experts completed the surveys independently and were instructed to consult the behavioral-cue rubric while scoring. Credibility (Q1) assessed whether a gene warranted inclusion on an AD target-discovery shortlist, based on the displayed evidence, prior knowledge, or both. Novelty (Q4) assessed its unfamiliarity as an AD therapeutic target based on the evaluator’s personal knowledge of the field. Both dimensions used five-point scales.

Primary comparisons used mean Q1 scores within each candidate group in each of the 16 questionnaires. The all-retained mean pooled ratings for Agora-listed and non-Agora retained genes. Group differences were calculated within questionnaires and averaged across questionnaires. The 95% confidence intervals for mean differences used Student’s *t* distribution with 15 degrees of freedom; *P* values were obtained from exact two-sided sign tests. As an expert-level sensitivity analysis, ratings from questionnaires completed by the same expert were pooled within each candidate group before calculating group means and paired differences. Each of the 14 experts therefore contributed one difference per comparison, with equal weight across experts. Confidence intervals and exact two-sided sign tests were recalculated using these expert-level differences, with 13 degrees of freedom for the Student’s *t* intervals. Rating distributions were calculated from individual ratings, whereas gene-level summaries averaged the ratings for each gene. Literature coverage *N_g_* was defined as the number of corpus publications containing a case-insensitive match to gene *g*’s symbol in the title or abstract, counting each publication once per gene. Matches immediately adjacent to letters, digits, or hyphens were excluded, and aliases were not expanded. Ordinary least-squares regressions of gene-level mean credibility and novelty on log_10_(*N_g_* + 1) were fitted separately for Agora-listed and non-Agora retained genes; rejected comparators were excluded from the fits. Exploratory analyses stratified retained genes by literature coverage (*N_g_ <* 100 or *N_g_ ≥* 100), reporting mean credibility ± s.e.m. across genes within each registry-by-coverage group.

